# ACTION ENHANCEMENT DRIVES THE EMBODIMENT OF A SUPERNUMERARY ROBOTIC DIGIT

**DOI:** 10.1101/2025.08.01.668241

**Authors:** John J Sykes, Francesca Genovese, Elena Mussini, Martina Fanghella, Guido Barchiesi, Domenico Prattichizzo, Simone Rossi, Corrado Sinigaglia

## Abstract

Wearable robotic supernumerary limbs enhance motor capabilities by leveraging principles from both engineering and neuroscience. Despite extensive research, how supernumerary limbs become embodied and the relationship between embodiment and artificially enhanced action possibilities remain largely unexplored. This gap has implications for device design and clinical adoption, as embodiment likely promotes patients’ acceptance and daily use of their supernumerary limb. Using an adapted proprioceptive drift paradigm from rubber hand illusion research, we investigated the embodiment of a robotic Soft Sixth Finger (SSF). Participants performed grasp-to-lift actions versus mere lift actions without object interaction, wearing the SSF on two different configurations: the palm and back of the hand. The Palm condition involved achievable actions without the SSF, while the Dorsal condition enabled otherwise anatomically impossible actions, extending beyond the physiological motor repertoire. Hand proprioceptive drift was measured before and after each task. Results showed that the SSF impacted body representation only when dorsally mounted. Proprioceptive drift was significantly higher for grasp-to-lift actions in Dorsal versus Palm conditions and higher for grasp-to-lift versus mere lift in the Dorsal condition only. This proves that SSF embodiment relates to extending, rather than substituting, users’ action possibilities. Future research must integrate behavioral, cognitive, and neural measurements across healthy and patient populations. Our study provides an ideal model for clinical application, demonstrating how enabling otherwise impossible actions drives robotic supernumerary finger embodiment.

## INTRODUCTION

The rapidly evolving field of wearable robotics (WR) combines principles from engineering and neuroscience to rehabilitate or enhance human action capabilities (Wu et al., 2014; Prattichizzo et al., 2021; Rossi et al., 2021; Eden et al., 2022). WR either augments the healthy body by artificially enabling actions beyond natural limits (e.g., in strength or workspace) (Eden et al., 2022) or serves a compensatory role by restoring motor function in individuals with impairments such as paresis by physically substituting a non-functional body part (Hussain et al., 2016; Dominjianni et al., 2021; Rossi et al., 2021; Umezawa et al., 2022).

The Soft Sixth Finger (SSF) is a wearable robotic device attached to the wrist that restores grasping abilities by affixing an underactuated supernumerary digit to the user’s hand (Hussain et al., 2016, 2017; Rossi et al., 2021). Even if the agent has impaired control over their fingers, the SSF enables basic grasp, lift, and object manipulation. This makes the SSF a potentially invaluable aid for individuals with hand paresis who can regain these abilities. Indeed, post-stroke patients have already given positive assessments of the SSF (Salvietti et al., 2021).

Despite extensive research on SSF and other WR devices from behavioral psychology (Wu et al., 2014; Hussain et al., 2015, Hussain et al., 2016; Meraz et al., 2018; Shafti et al., 2021; Arai et al., 2022; Fasoulas et al., 2024) and cognitive neuroscience (Hussain et al., 2017; Kieliba et al., 2021; Rossi et al., 2021; Liu et al., 2024; Katmah et al., 2024), fundamental questions about user-device relationships remain unanswered. Perhaps most critically: do users embody these devices, and if so, how does this embodiment process occur?

The challenge becomes particularly complex when considering that WR devices often enable actions only possible while wearing the device, which falls outside the usual motor repertoire, raising questions about whether these devices truly become incorporated into users’ body schema. How WR devices can be embodied and how this embodiment relates to enhanced action possibilities remains largely unexplored. However, such questions have significant implications for WR’s future development and implementation. The present study aimed to address this gap by taking a fundamental first step toward clarifying how SSF embodiment links to enhanced user action possibilities.

Enhancing action possibilities can be achieved in two ways (Eden et al., 2022). The first works by improving pre-existing action performance, i.e., by enhancing strength, stability, precision, or workspace. For instance, Shafti et al. (2021) utilized a robotic third thumb to expand the agent’s workspace when playing the piano. They found that, after approximately 40 minutes of use, novice and expert piano players could incorporate the device to reach more piano keys while playing. The second enhancement mode pertains to actions otherwise absent from the users’ motor repertoire, a comparatively underexplored factor in the literature (but see Kieliba et al., 2021; Umezawa et al., 2022).

We investigated this second mode of action enhancement by exploiting the adaptable placement of the SSF and comparing two distinct configurations during a grasp-to-lift task. Despite working with healthy participants who possessed intact natural grasping abilities, we manipulated the functional nature of their actions by mounting the SSF with two different configurations. In the Palm condition, participants wore the SSF on the wrist with the palm side facing outward, while in the Dorsal condition, the SSF was positioned on the back of the hand. The main task involved repeatedly grasping and lifting a three-dimensional object. In a control condition, participants activated the device by opening and closing the SSF without targeting any object.

The two SSF configurations created fundamentally different action modalities. In the Palm condition, participants performed actions that remained functionally achievable without the SSF by using their natural thumb. By contrast, the Dorsal configuration enabled participants to execute grasp-to-lift movements that would be anatomically impossible without SFF. This dorsal placement temporarily endowed participants with novel action possibilities that depended entirely on the use of SSF.

We employed proprioceptive drift to measure SSF embodiment. This measure has been reliably used to assess the embodiment of external entities, such as rubber hands (Botvinick & Cohen, 1998; Tsakiris & Haggard, 2005; Capelari et al., 2009; Gallagher et al., 2021). Moreover, proprioceptive drift has been demonstrated to be sensitive to novel action possibilities with respect to virtual or robotic hands (Kammers et al., 2009; Kalckert & Ehrsson, 2012; Romano et al., 2015; Shibuya et al., 2017).

Proprioceptive drift was measured before and after participants completed both the grasp-to-lift actions and the empty-handed lift movements, while wearing the SSF on the palm or the back of the hand. We quantified embodiment using proprioceptive drift along the horizontal plane in the direction of SSF placement.

We compared the proprioceptive drift between object-directed grasping actions and simple device activation movements to evaluate the influence of action enhancement. Our hypothesis predicts that if SSF embodiment relates to action enhancement, proprioceptive drift will be modulated by hand position, with effects present in the Dorsal but not the Palm condition. This prediction stems from the fundamental difference between conditions. While the palm-mounted SSF replicated thumb functionality without significantly expanding participants’ motor repertoire, the dorsal-mounted SSF enabled grasping actions that extended their physiological possibilities for action. The SSF would be embodied, impacting body representation, as it enables new action possibilities, allowing users to perform actions that would otherwise be impossible or no longer possible.

## MATERIALS AND METHODS

### Power analysis

Since we utilize a classic rubber hand illusion measure to determine the sample size of our experiment, we refer to Kalckert & Ehrsson (2012), as they employed the rubber hand illusion and the proprioceptive drift measure in a similar experimental design (active and passive, with congruent and incongruent conditions). The power analysis results, performed using G^*^power (Faul et al., 2007), indicate that a total sample size of 7 participants is required (effect size f = 0.5, alpha = 0.05, power = 0.80, 2 positions x 2 conditions). To detect a medium effect size (0.25), 24 participants were necessary; however, since this measure has never been used to assess proprioceptive drift after using the sixth soft finger, we employed 32 participants.

### Participants

We collected data from 32 participants (16 women, aged 19-53, mean age 22.13, 6 of whom were left-handed, with no neurological disorders and vision either corrected or normal). Participants were recruited from the University of Milan and received course credits. The study complied with the revised Helsinki Declaration (World Medical Association General Assembly, 2008) and was approved by the local Ethical Committee. All participants provided written informed consent for their participation and completed the Edinburgh Handedness Inventory questionnaire to determine which hand the SSF was placed in.

### Soft Sixth Finger (SSF)

The Soft Sixth Finger (SSF) is a wearable supernumerary robotic device developed to restore grasping capacities in those suffering from paresis by providing a modular, underactuated artificial digit attached to the user’s wrist (Prattichizzo et al., 2014; Hussain et al., 2016). This robotic digit is capable of flexion and extension and can be activated as needed through various interface modalities. In our version, these movements are controlled by two buttons on an elastic ring worn on the finger of the other hand. The SSF offers a low-complexity yet highly wearable solution for grasping tasks. It can reliably grasp objects of various shapes and sizes due to its intrinsically compliant structure, independent of the subject’s skeletal structure. Objects are gripped between the digit and the hand or forearm. Subjects control the onset of the action and can reverse this movement, but the SSF moves essentially autonomously and wraps around objects automatically before stopping. The digit is mechatronically composed of a tendon-driven system connected by a thread and powered by a single motor. The main body of the SSF is attached to the user’s wrist with a velcro strap, allowing it to be placed in more than one position (i.e., underneath and over the wrist). The SSF includes an electronic board taped to the same arm as the device. The whole system weighs about 160 grams.

### Experimental Structure

Participants were asked to perform a sequence of actions while wearing the SSF. The potential embodiment of the SSF was assessed by measuring their proprioceptive drift before and immediately after the action performance. The proprioceptive drift was measured while participants were not wearing the SSF.

### SSF Action Tasks

Based on the Edinburgh handedness inventory questionnaire results, the SSF was placed on the participant’s dominant hand. The SSF experiment included two experimental conditions: Palm and Dorsal. In the Palm condition, participants wore the SSF on the palm side, while in the Dorsal condition, the SSF was positioned on the back of the hand (Figure 1).

**Figure 1.**
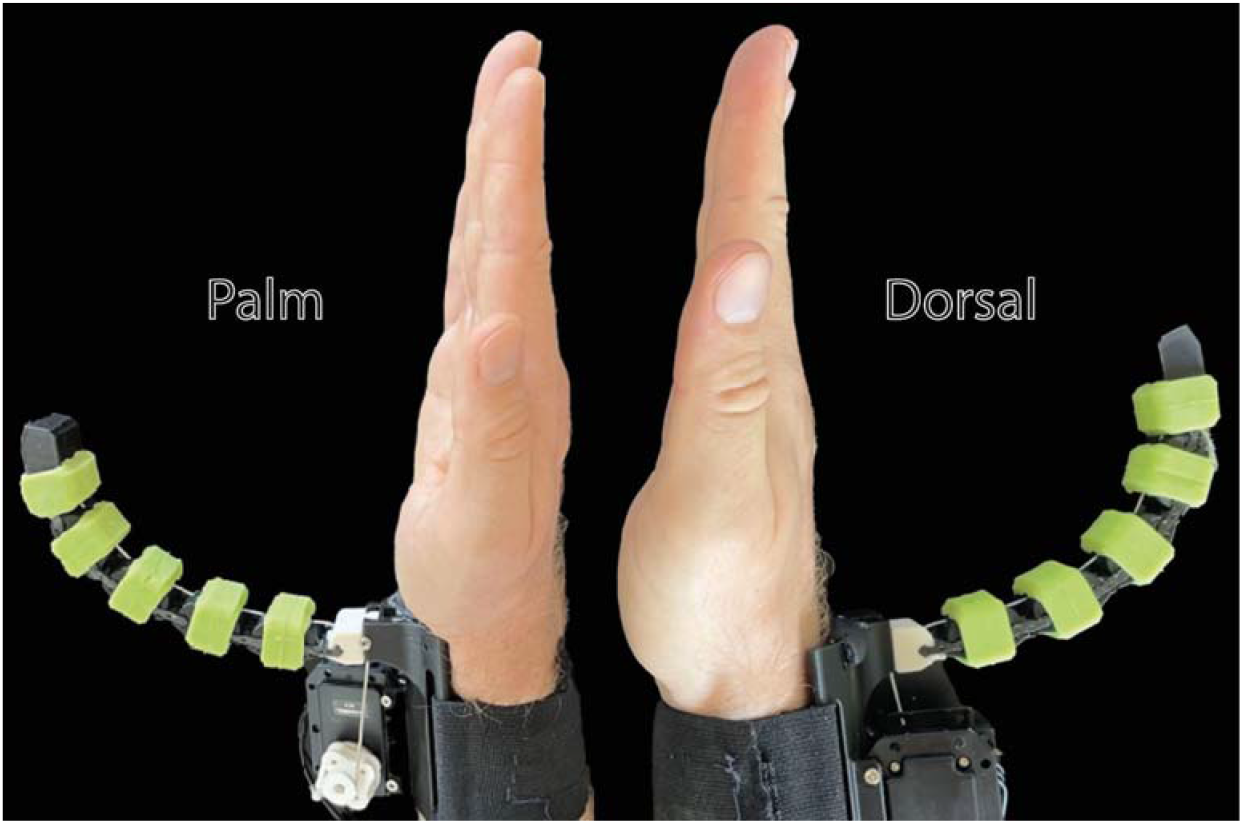
The two experimental conditions. On the left, the Palm condition: the SSF is mounted on the palm. On the right, the Dorsal condition: the SSF is mounted on the back side of the hand.

For each condition, participants performed a Real and a Control task (Figure 2). In the Real Task, participants lifted an empty aluminum bottle (22 cm tall; 94g weight) approximately 15 cm, using only the SSF to grasp it while keeping their biological fingers stationary. Afterward, the bottle was placed back on the table, the SSF opened, and the arm moved back slightly (Two illustrative video clips are enclosed as supplementary movies). In the Control task, subjects had to open the SSF, raise their arm approximately 15 cm, lower it, and then close the SSF without moving any biological fingers and interacting with any object (Two illustrative video clips are enclosed as supplementary movies). Participants performed both tasks with closed eyes, allowing them to rely solely on tactile and auditory feedback related to finger activation and closure during the grasp-to-lift actions (Real task) or the empty-handed lift movements (Control task).

**Figure 2.**
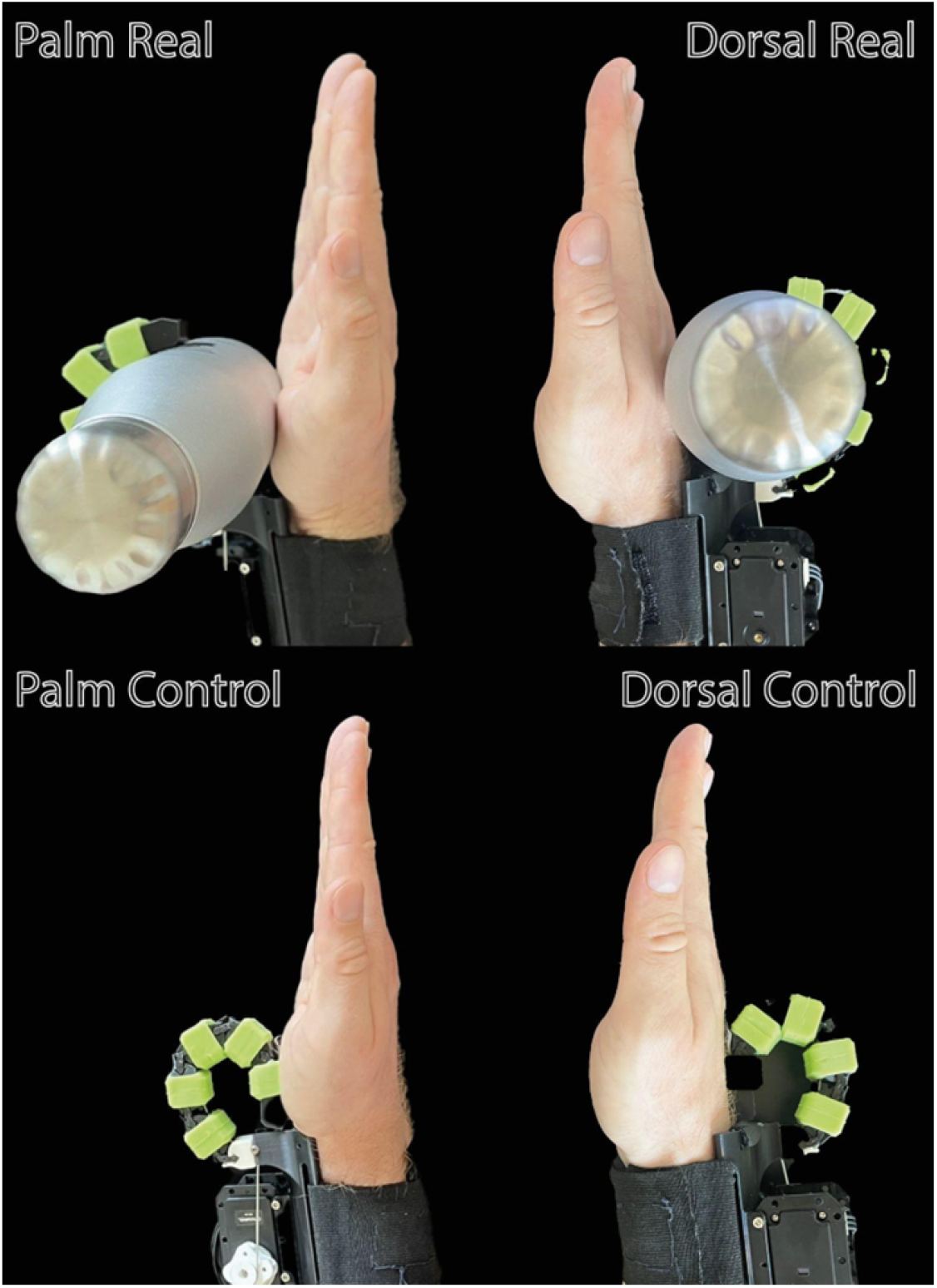
The two tasks (Real and Control) under the two experimental conditions (Palm and Dorsal). In the Real task, participants grasped and lifted an empty bottle, while in the Control task, they lifted and closed the sixth finger without interacting with any object. Illustrative video clips are enclosed as supplementary movies.

The experiment consisted of two blocks. In each block, participants performed the two tasks (Real and Control) while wearing the SSF on either the palm (Palm condition) or the back (Dorsal condition) side (e.g., block 1: PalmReal and PalmControl; block 2: DorsalReal and DorsalControl). Each block’s combination of tasks and conditions was randomly counterbalanced across participants. Each task lasted 5 minutes, and on average, they produced the same number of movements in the Real and Control tasks. Subjects underwent a 5-minute break between the Real and Control tasks and a 10 minute break between the first and second blocks. Before each block, participants completed a 5-minute familiarization session with the SSF.

### Proprioceptive Drift

We adapted the proprioceptive drift measure (Kalckert & Ehrsson, 2012). Based on the Edinburgh handedness inventory questionnaire results, a fine-tipped felt-tip pen was tied to the participants’ index finger of the non-dominant hand. Participants placed their hands in a vertical position with respect to the table to better match the hand’s position when using the SSF.

To measure participants’ proprioceptive drift, subjects indicated ten times on an A4 sheet of paper where they felt their concealed hand was. A sheet of A4 graph paper (28 cm) was placed on top of a small, breakfast-like table, which was positioned on the larger table where participants sat. Participants put their dominant hand (i.e., the hand that used the SSF) in an indicated position underneath the small table, behind a cloth that concealed it. Participants were first instructed to close their eyes. Subsequently, using a pen tied to the index finger of their non-dominant hand, they indicated on the graph paper where they felt their hand was, using their thumb as a point of reference (Figure 3). Subjects made a total of 10 indications on the A4 sheet immediately after an experimenter had counted to a corresponding number. The proprioceptive drifts were measured 6 times. Twice as a baseline before each block (e.g., Palm/Baseline and Dorsal/Baseline), and once after each condition’s and task’s completion (e.g., PalmReal, PalmControl, DorsalReal, DorsalControl). SSF familiarization preceded each post-baseline block

**Figure 3.**
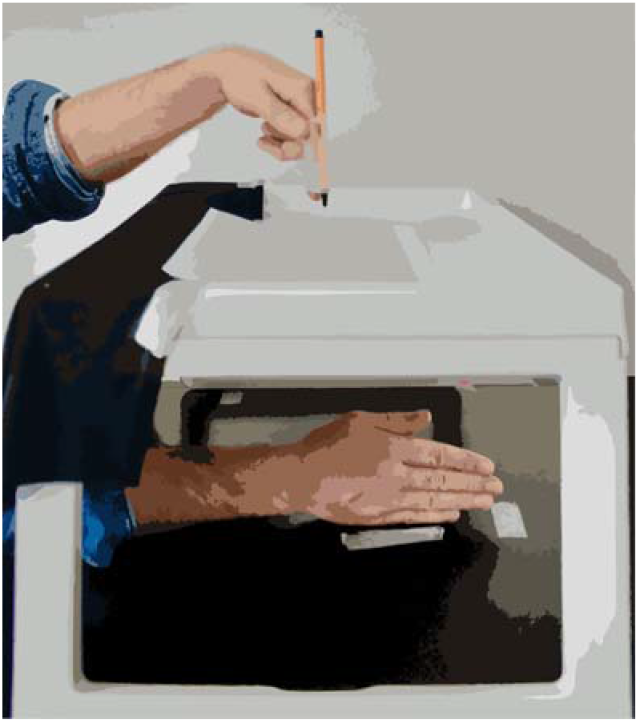
The figure shows the proprioceptive drift measure’s setting: participants place their dominant hand under the table within a delimited space and, with the other hand, indicate ten times where they feel their hand is, using the thumb and reference, pointing a pen on a sheet of graph paper (28 cm) placed on the table.

## Data Analysis

### Proprioceptive Drift Index

We calculated the proprioceptive drift index for each participant by subtracting the mean of the ten baseline points from the mean of the ten points in each condition for each task. We thus obtained two values for the Palm condition and two for the Dorsal condition (PalmReal - Baseline, PalmControl - Baseline, DorsalReal - Baseline, DorsalControl - Baseline). In the Dorsal condition, positive values represent a rightward shift towards the position of one’s thumb, as identified in the baseline. In contrast, in the Palm condition, negative values represent this shift. Since left-handed participants were in the opposite direction, the sign of the proprioceptive drift index of each condition was inverted. To assess the impact of action enhancement on SSF embodiment as measured by the proprioceptive drift index, we performed a two-way repeated-measure 2×2 ANOVA with CONDITION (Palm and Dorsal) and TASK (Real and Control) as within-subject factors.

## RESULTS

The two-way ANOVA (CONDITION x TASK) produced a significant main effect of CONDITION (F_1,31_ = 9.18, p = **0.005**, partial-η^2^ = 0.229) and a significant interaction of CONDITION X TASK (F_1,31_ = 10.93, p = **0.002**, partial-η^2^ = 0.261) (Figure 4; Table 1). We performed t-tests between Dorsal and Palm in the Real task and found a significant effect with Dorsal Real higher than Palm Real (t_31_ = 4.071, p < **.001**, Cohen’d = 0.72). No differences were found between Dorsal and Palm in the Control task (t_31_ = 1.293, p = 0.206, Cohen’d =0.229). We also performed t-tests (Bonferroni corrected for four comparisons, p < 0.0125) between Real and Control within each Condition, and we found a significant effect for Dorsal with the Real task higher than Control (t_31_ = 3.276, p = **0.003**, Cohen’d = 0.579). No differences were found between the Real and Control tasks for the Palm condition (t_31_ = -0.968, p = 0.341, Cohen’d = -0.171). (Table 2)

**Table 1.**

| Anova Repeated Measures 2*2 |  |  |  |  |  |  |
| --- | --- | --- | --- | --- | --- | --- |
| Factors | Sum of Squares | df | Mean Square | F | p | $\eta^2p$ |
| Conditions | 57.352 | 1 | 57.352 | 9.182 | **0.005** | 0.229 |
| Residuals | 193.630 | 31 | 6.246 |  |  |  |
| Task | 5.064 | 1 | 5.064 | 2.397 | 0.132 | 0.072 |
| Residuals | 65.499 | 31 | 2.113 |  |  |  |
| Conditions × Task | 16.560 | 1 | 16.560 | 10.933 | **0.002** | 0.261 |
| Residuals | 46.956 | 31 | 1.515 |  |  |  |

**Table 2.**

| Paired Samples T-Test |  |  |  |  |  |
| --- | --- | --- | --- | --- | --- |
| Measure 1 | Measure 2 | t | df | P-value<br>($p < 0.0125$) | Cohen's d |
| DorsalReal-Baseline | PalmReal-Baseline | 4.071 | 31 | **< .001** | 0.72 |
| DorsalControl-Baseline | PalmControl-Baseline | 1.293 | 31 | 0.206 | 0.229 |
| DorsalReal-Baseline | DorsalControl-Baseline | 3.276 | 31 | **0.003** | 0.579 |
| PalmReal-Baseline | PalmControl-Baseline | -0.968 | 31 | 0.341 | -0.171 |

**Figure 4.**
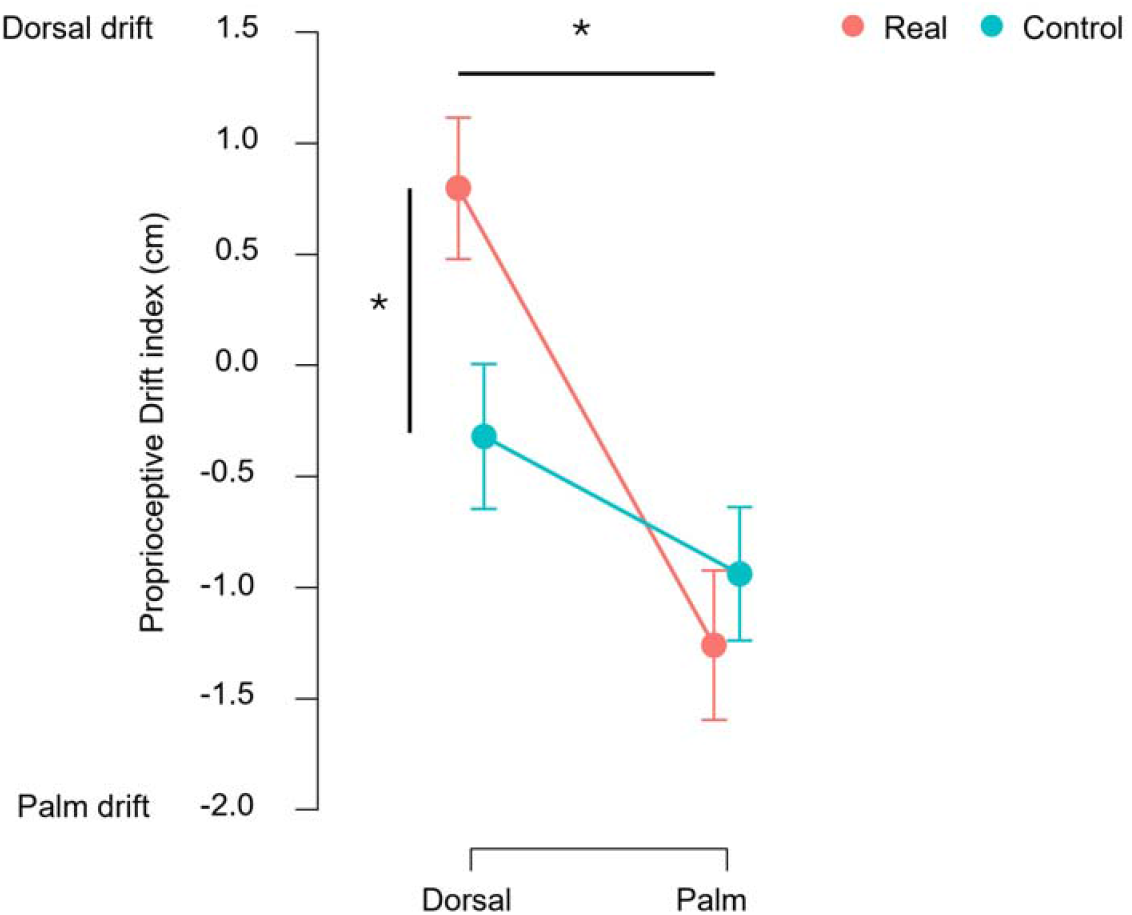
The plot shows the subtraction of the mean of the ten points of the baseline from the mean of the ten points of each condition for each task. This way, we calculated the shifting in the perception of one’s hand position from the baseline (i.e., proprioceptive drift index represented by the dots). Error bars represent standard error. Since the hand was placed vertically during the measurement, negative values represent a shift towards the palm, while positive values represent a shift towards the dorsal side. The “^*^” symbol with the horizontal line signals a significant difference in the proprioceptive drift index between the Dorsal and Palm conditions in the Real task. The “^*^” symbol with the vertical line signals a significant difference in the proprioceptive drift index between the Real and Control tasks in the Dorsal condition (i.e., the significant interaction between Condition x Task).

## DISCUSSION

The current study aimed to investigate how the use of a wearable robotic device, such as the Soft Sixth Finger, can impact body representation and how this impact relates to enhancing action possibilities. We compared grasp-to-lift actions with mere, empty-handed lift actions, performed without grasping any object, with the SSF positioned on either the palm or the back of the hand. To assess the impact of action enhancement on body representation, we measured proprioceptive drift before and after participants performed both grasp-to-lift and empty-handed lift actions while wearing the SSF on the palm or the back of the hand.

Results show that the SSF impacted body representation when mounted on the back of the hand but not on the palm. Specifically, proprioceptive drift was significantly higher for grasp-to-lift actions in the Dorsal than in the Palm condition. Furthermore, proprioceptive drift was greater for grasp-to-lift actions than simple lift actions in the Dorsal condition only. The difference between tasks (grasp-to-lift vs. empty-handed lift) in the Dorsal condition indicates that the effect cannot be attributed solely to SSF placement on the back of the hand. Similarly, the difference between conditions (Dorsal vs. Palm) during both grasp-to-lift tasks indicates that the effect cannot be due to the object’s weight, as the same object (empty bottle) was grasped in both conditions but only produced drift in the Dorsal condition.

The present findings suggest that agents can embody a WR device, such as the SSF, to the extent that it endows their body with novel action possibilities. When the SSF was mounted on the back of the participant’s hand, the sensed location of the participant’s hand drifted toward that of the acting device. The fact that embodiment occurred uniquely in the Dorsal condition likely stems not from the SSF’s placement per se but from the unique type of action it enabled. In the palm configuration, participants could already perform functionally similar actions with their natural hand. In contrast, the dorsal configuration enabled biomechanically impossible actions without the SSF, thereby classifying it as an effective action enhancement because it extends action possibilities rather than merely implementing pre-existing actions. Because specific action extensions are uniquely possible whilst wearing the SSF, participants appeared more likely to embody it.

As a limitation of the study, we did not examine the neural representations that may underlie this embodiment process, which was clearly evident at the behavioral level. However, previous research has demonstrated that the active use of robotic digits critically impacts body representation. Segura Meraz et al. (2018) studied participants using a robotic thumb controlled by the contralateral thumb to touch indicated fingers, with electrostimulation feedback sequentially. Results showed progressive incorporation of the robotic finger and modification of the controlling finger’s representation via proprioceptive drift. Similarly, Umezawa et al. (2022) examined a robotic sixth finger during bending and tapping tasks. While no significant proprioceptive drift occurred, questionnaires indicated embodiment and changes in finger localization correlated with ownership reports. Kieliba et al. (2021) trained participants to perform bimanual tasks (e.g., holding and stirring) using one hand augmented with a robotic thumb. Training altered natural finger coordination and transiently reduced neural hand representation in the sensorimotor cortex. Rossi et al. (2021) showed the immediate formation of new bioartificial corticospinal synergies for hand muscles using transcranial magnetic stimulation (TMS) during motor imagery tasks after training with a supernumerary thumb.

The novelty of our study lies in demonstrating how action enhancement drives the embodiment of supernumerary robotic digits. Ongoing debates question the feasibility of robotic body augmentation (Aoyama et al., 2019; Makin et al., 2020; Dominijianni et al., 2021; Prattichizzo et al., 2021; Prattichizzo et al., 2014; Arai et al., 2022), with some arguing that anatomical, neural, or cognitive constraints limit integration of new WR functions into body representation (Makin et al., 2020; Dominijianni et al., 2021; Marucci et al., 2024; De Vignemont, 2024). Our findings suggest that enabling previously impossible actions due to hand-related biomechanical constraints induces the embodiment of a supernumerary digit. This effect was absent when the SSF was palm-mounted, likely because grasp-to-lift actions did not alter the hand’s functional representation. Although the SSF operates independently of natural fingers, these actions remained achievable using the biological fingers alone. However, this does not preclude the embodiment of a palm-mounted SSF if its active use genuinely expands users’ action possibilities.

Unlike previous studies requiring 10-15 minutes (Kieliba et al., 2021) or 40 minutes (Shafti et al., 2021), only 5 minutes of performing grasp-to-lift actions were sufficient to induce SSF embodiment in our study, as measured by proprioceptive drift. This timing aligns with Romano et al. (2015), who found that 5 minutes induced embodiment effects in a robotic hand setup mirroring the rubber hand illusion. The rapid embodiment may reflect the ease of skill acquisition due to the SSF’s independent functioning (Eden et al., 2022). Although participants performed a novel action, they successfully completed grasp-to-lift tasks after only 5 minutes of familiarization, with minimal errors. Furthermore, visual deprivation may have facilitated embodiment during tasks, with haptic feedback and motor acoustic cues playing critical roles in integrating this novel, unseen action into body representation (Dadarlat et al., 2015).

If confirmed, our findings have both theoretical value and significant implications for SSF and, more generally, WR development and implementation, as embodiment likely promotes device acceptance and daily use (Bidiss & Chau, 2007; Segura Meraz et al., 2018; Sato et al., 2018; Franco et al., 2021; Kieliba et al., 2021). The SSF was developed for patients with paresis (Hussain et al., 2016, 2017; Prattichizzo et al., 2014, 2021). We found that SSF embodiment occurs when the device enables actions that would otherwise be impossible. This principle may apply to biomechanically impossible actions and lost action capabilities due to a brain or spinal cord lesion. For patients with impaired grasping actions and preserved sensory pathways, artificially restoring these action possibilities via palm-positioned SSF may induce embodiment similar to that observed in healthy subjects with dorsal positioning. The proprioceptive drift index may then be used to assess SSF embodiment over time during the various phases of rehabilitation and recovery of hand functionality.

Our findings indicate that supernumerary devices become embodied only when they enhance action capabilities through learning new bioartificial actions. This principle should guide future research integrating behavioral and cognitive measurements while exploring the underlying neural mechanisms in healthy participants and patients. Our study provides an ideal model for clinical application, demonstrating how enabling otherwise impossible actions drives robotic supernumerary finger embodiment.

## Acknowledgments

This article was supported by the Department of Philosophy ‘Piero Martinetti’ of the University of Milan with the Project “Departments of Excellence 2023-2027” awarded by the Italian Ministry of Education, University and Research (MIUR) (to JS, FG, EM, MF, GB, and CS) and the PRIN 2022 grant “The extended hand: psychophysical and neural foundations of a robotic supernumerary finger’s use for grasping augmentation or recovery” (2022J72LFW_001/2 toJS, DP, SR and CS). We would like to thank Angelica Kaufmann for the figures and video clips.

## Author Contributions

Conceptualization, JJS, FG, and CS; Methodology, FG, JJS, and GB; Formal analysis, FG and GB; Investigation, JJS, FG, EM, and MF; Writing –Original Draft, CS, JJS, and FG; Writing – Review and Editing, CS, DP, and SR; Visualization, FG and CS; Supervision, CS, DP, and SR; Funding Acquisition, CS, DP and SR.

## Transparency and Openness

Data, Analysis code, and Research Materials can be found at https://osf.io/d9vt4/

